# The unique role of *nucS*-mediated non-canonical mismatch repair in *Mycobacterium tuberculosis* resistance evolution

**DOI:** 10.1101/2025.06.19.660520

**Authors:** Isabel Martín-Blecua, José Ramón Valverde, Pablo García-Bravo, Ángel Ruiz-Enamorado, Rafael Prados-Rosales, Jorge Sastre-Domínguez, Lahari Das, William R. Jacobs, Álvaro San Millán, Jesús Blázquez, Sonia Gullón

## Abstract

DNA surveillance mechanisms play a vital role in maintaining genome stability and minimizing mutation rates. One such mechanism, post-replicative mismatch repair (MMR), corrects replication errors that escape DNA polymerase proofreading activity. In most bacteria and eukaryotes, MMR is orchestrated by MutS and MutL proteins. However, certain archaeal and actinobacterial species, including the major human pathogen *Mycobacterium tuberculosis*, lack these components. Instead, they rely on the nuclease EndoMS/NucS, a structurally distinct enzyme that governs a non-canonical MMR pathway. Given that *M. tuberculosis* acquires drug resistance exclusively through chromosomal mutations, understanding mutation rate regulation in this pathogen is critical. Nevertheless, despite its anticipated significance, the role of NucS in drug resistance evolution remains largely unexplored in this organism.

This study investigates NucS function in *M. tuberculosis* and uncovers a unique resistance dynamic distinct from other Actinobacteria. While *nucS* deletion alters the mutational spectrum, it minimally affects the emergence of rifampicin-, isoniazid-, and ethambutol-resistant mutations, in stark contrast to its role in other Actinobacteria. We demonstrated that this atypical behaviour is not attributable to the presence of a single NucS polymorphism, R144S, in the NucS sequence of the *M. tuberculosis* reference strain H37Rv, which differs from the NucS consensus sequence. Constructing and analysing an H37Rv variant possessing the NucS consensus sequence revealed a subtly altered mutational spectrum but unchanged mutation rates. Notably, database analysis of the R144S polymorphism in clinical isolates revealed its prevalence and significant association with ethambutol resistance. These findings challenge the established view that *nucS* serves as a genome stability guardian that minimizes mutation rates in *M. tuberculosis*, suggesting additional mismatch repair mechanisms beyond NucS or a highly efficient replication system in this pathogen.

## Introduction

To maintain genome stability, cells use a number of DNA surveillance and correction processes. Among these processes, the fidelity of DNA replication is a key factor in keeping mutations at a low rate. This fidelity is ensured by both base selection and the 3’-5’ proofreading activity of replicative DNA polymerase, as well as the postreplicative mismatch repair (MMR). The contribution of each of these activities to the error rate is roughly estimated at 10^-5^, 10^-2^, and 10^-3^, respectively, accounting for an overall error rate of 10^-10^ (1). MMR repairs errors (mismatches) that escape the correcting activity of DNA polymerase, relying on proteins of the MutS and MutL families for function (2). However, certain archaeal and most actinobacterial species, including the causative agent of tuberculosis (TB) *Mycobacterium tuberculosis*, lack the *mutSL* genes yet exhibit low mutation rates, suggesting the existence of an alternative MMR system (3). The EndoMS/NucS protein (hereafter referred to as NucS for simplicity) plays a key role in this alternative MMR pathway in these prokaryotes (3–5). Initially identified in the archaeal species *Pyrococcus abyssi*, NucS is a two-domain protein with an N-terminal DNA-binding domain connected by a short linker to the C-terminal catalytic domain, defining a new family of structure-specific DNA endonucleases(6). Further studies characterized its biochemical activity, revealing that archaeal NucS is an endonuclease that recognizes mismatched sites arising during DNA replication. Once a mismatch is recognized, NucS introduces double-strand breaks (DSBs) flanking the site of the mismatch, generating 5’sticky ends with five overhanging nucleotides leaving the mismatched base in the middle. The cleavage activity of the endonuclease NucS was shown to be enhanced by the sliding clamp (PCNA in Archaea and β-clamp in Bacteria) (7, 8).The interaction between NucS and β-clamp occurs through a sequence of five amino acid residues present at the end of the C-terminal NucS sequence. These DBSs must be processed to restore the correct base pair, although the specific repair mechanism remains undefined. Recent studies suggest that neither homologous recombination (HR) nor non-homologous end joining participate in this pathway in *Mycobacterium smegmatis*(9, 10).

The first in vivo evidence of NucS-mediated MMR activity in Actinobacteria was provided by Castañeda-García et al in 2017 (3), who identified *nucS* (*MSMEG_4923*) as an antimutator gene in *M. smegmatis* through a transposon mutagenesis screen for mutants with high rates of spontaneous rifampicin resistance. The deduced amino acid sequence of this gene had a 27% sequence identity with NucS from *P. abyssi*. Genetic and biological analysis showed that the *nucS*-null (Δ*nucS*) variant of *M. smegmatis* exhibited dramatically increased mutation rates, a transition-biased mutational spectrum and elevated recombination rates between non-identical (homeologous) DNA sequences, phenotypes almost identical to those produced by the MMR deficiency in other organisms relying on MutS and MutL activity (3). The hypermutator phenotype of actinobacterial *nucS*-null mutants was confirmed in *Streptomyces coelicolor* (3) and further verified in *Corynebacterium glutamicum* (4, 11), *Mycobacterium abscessus* (12) and *Streptomyces ambofaciens* (13). Purified NucS from *C. glutamicum*(4, 11) and *S. ambofaciens* (13) demonstrated enzymatic activity similar to their archaeal counterparts, efficiently correcting transition mutations. NucS was found to efficiently correct mainly transition mutations (A:T>G:C and G:C>AT) in Actinobacteria *in vivo*, as deduced from analysis of mutations conferring antibiotic resistance(3, 4) and mutation accumulation (MA) experiments(11, 14). Fluctuation tests or estimation of mutant frequency of antibiotic resistant mutants showed that deletion of *nucS* conferred a hypermutable phenotype across all tested Actinobacteria(3, 4, 11–13), increasing mutation rate by two orders of magnitude compared to wild-type strains. This underscores the critical role of NucS in regulating genome stability and contributing to the evolution and adaptability of these clinically and industrially important species.

*M. tuberculosis* remains one of the deadliest pathogens in human history, causing tuberculosis (TB). TB has likely regained its status as the world’s leading infectious disease killer, with more than 10.8 million cases and 1.25 million deaths in 2023 (15). The rise of antibiotic resistance in *M. tuberculosis* has worsened this public health challenge, complicating treatment protocols and undermining global TB control efforts worldwide. Multidrug resistant strains (MDR), some of them resistant to most or all effective antibiotics, have emerged steadily over decades (16, 17). Recently, rifampicin resistant (RIF-R) *M. tuberculosis* has been included in the 2024 WHO list of critical bacterial priority pathogens (18). Unlike most bacterial pathogens that acquire drug resistances mainly by horizontal gene transfer, *M. tuberculosis* develops drug resistance exclusively through chromosomal mutations(19), emphasizing the importance of understanding mutation rate regulation in this species (20). There is compelling evidence that hypermutable strains, often associated with defects in MMR components, play a significant role on the development of antibiotic resistance, virulence, persistence and transmissibility in chronic bacterial pathogens, such as *Pseudomonas aeruginosa* in cystic fibrosis patients (21) (22). Even small increases in mutation rates can have substantial effects on the evolution of antibiotic resistance (20, 23, 24). However, the role of *nucS* in regulating the development of antibiotic resistance in *M. tuberculosis* has not been explored. To investigate this, we have constructed and analysed a Δ*nucS* mutant and its corresponding *nucS*-complemented variant in the *M. tuberculosis* H37Rv model strain. Using fluctuation tests, we assessed the rate at which *M. tuberculosis* acquires resistance to the first-line antitubercular drugs rifampicin (RIF), isoniazid (INH) and ethambutol (EMB). Furthermore, as H37Rv harbours a unique polymorphism (R144S) relative to the *M. tuberculosis* consensus NucS sequence (25), we constructed an engineered variant of H37Rv harbouring the NucS consensus sequence and analysed how it affected the mutational spectrum and resistance rates. Finally, prevalence of R144S polymorphism among clinical *M. tuberculosis* genomes and its association with drug resistance has been analysed.

## Results

### Deletion of nucS does not affect the rate of RIF-R spontaneous mutations

To assess the impact of NucS on mutation rates in *M. tuberculosis*, we constructed a Δ*nucS* derivative in the *M. tuberculosis* strain mc^2^6230, an attenuated variant of the model strain H37Rv(26). Deletion of *nucS* did not significantly increase the rate of RIF-R spontaneous mutations (significance defined by nonoverlapping 99% CI). Specifically, the rates of spontaneous RIF-R mutants were 5.49 ×10^-9^ (95% CI: 4.65-6.63 ×10^-9^) for wild-type (WT) and 1.55×10^-8^ (95% CI: 1.24-1.89 ×10^-8^) for the Δ*nucS* mutant (Fig. 1A and Supplementary Table 1). This represented a mere 2.83-fold increase upon *nucS* deletion. These results were highly unexpected, given that deletion of *nucS* in all tested Actinobacteria, including *M. smegmatis*(3), *S. coelicolor*(3), *S. ambofaciens*(13) and *C. glutamicum*(4, 11, 27), consistently led to a two-order magnitude increase in spontaneous RIF-R mutation rates (Fig. 1B).

**Figure 1.**
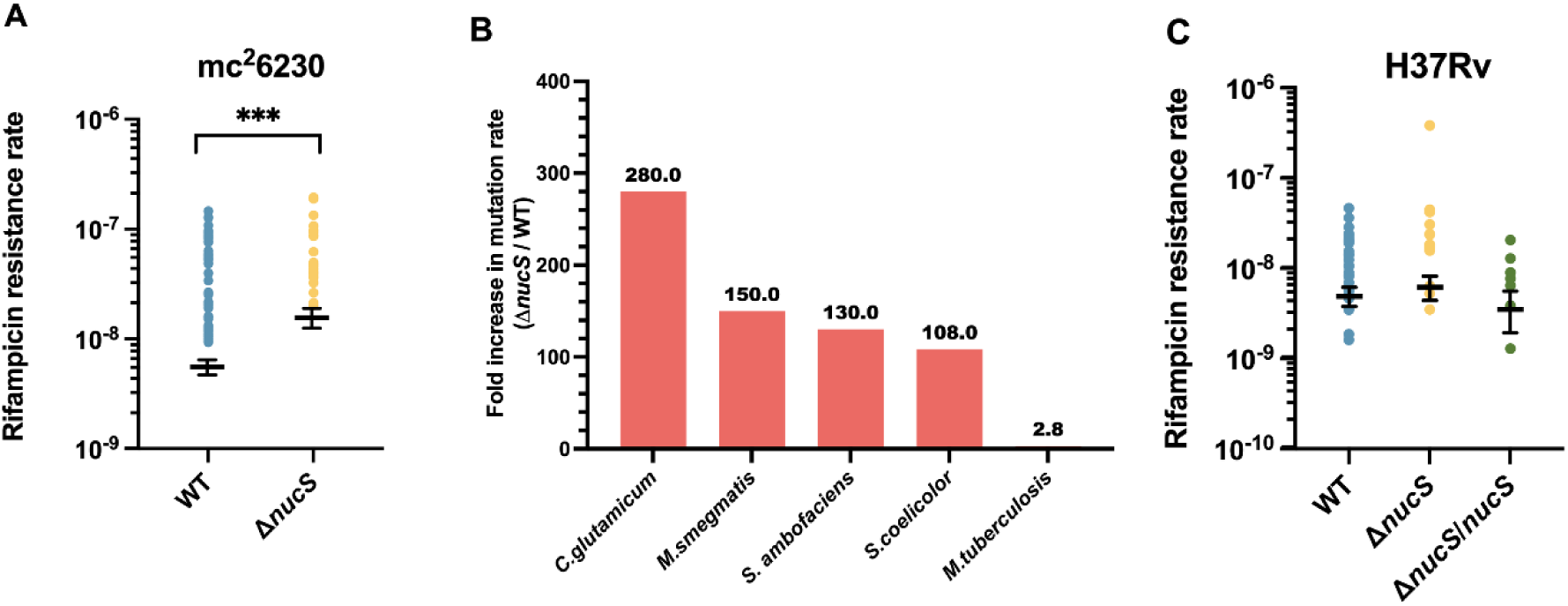
Effect of *nucS* deletion on the rate of spontaneous RIF-R mutants. Mutation rate for RIF-R spontaneous mutants in strain mc^2^6230 and its Δ*nucS* derivative (A). Fold increase for RIF-R in mutation rate for Δ*nucS* derivatives of *M. smegmatis*(3), *S. coelicolor*(3), *S. ambofaciens*(13), *C. glutamicum*(4, 11, 27) and *M. tuberculosis* strain H37Rv mc^2^6230 (this work) (B). RIF-R mutation rates of strains H37Rv and their derivatives Δ*nucS* and Δ*nucS*/*nucS* (C). Differences are not statistically significant in C.

To determine whether this non-mutator phenotype was specific to the H37Rv avirulent derivative, we generated a Δ*nucS* mutant in the virulent parental strain H37Rv and analysed RIF-R mutation rates in wild-type, Δ*nucS* (H37Rv Δ*nucS*), and its *nucS*-complemented derivative (H37Rv Δ*nucS*/*nucS*). Again, the Δ*nucS* strain exhibited a RIF-R mutation rate comparable to the wild-type strain. Specifically, the rates of spontaneous RIF-R mutants were 4.66 ×10^-9^ (95% CI: 3.58-5.86 ×10^-9^), 5.88 ×10^-9^ (95% CI: 4.18-7.79 ×10^-9^) and 3.33 ×10^-9^ (95% CI: 1.82-5.29 ×10^-9^) for the wild-type, Δ*nucS,* and Δ*nucS*/*nucS* variants, respectively (Fig 1C and Supplementary Table 1). Similar results with the H37Rv Δ*nucS* strain were observed by K. Murphy and C. Sassetti (personal communication). These findings challenged the hypothesis that *nucS* serves as a guardian of genome stability in *M. tuberculosis,* strongly suggesting that, unlike in other Actinobacteria, *nucS* does not modulate mutation rates in this species.

### Effect of nucS-deletion on the rate of INH-R and EMB-R spontaneous mutations

To further evaluate the role of NucS in modulating mutation rates in *M. tuberculosis*, we tested the mutator phenotype using two additional first-line antitubercular drugs: isoniazid and ethambutol. The spontaneous INH-R mutation rates were 8.09 ×10^-7^ (95% CI: 6.95-9.20 ×10^-7^), 1.57 ×10^-6^ (95% CI: 1.37-1.76 ×10^-6^) and 9.96 ×10^-7^ (95% CI: 7.89-11.9 ×10^-7^) for WT, Δ*nucS* and *nucS*-complemented H37Rv strains, respectively (Fig. 2A and Supplementary Table 1). This represents a modest 1.94-fold increase in mutation rate upon *nucS*-deletion, with restoration of near WT levels upon complementation. Similarly, the rates of spontaneous EMB-R mutations were 9.57 x 10^-9^ (95% CI: 6.20-13.30×10^-9^), 6.88 x 10^-9^ (95% CI: 4.10-10.40 x 10^-9^) and 1.42 x 10^-9^ (95% CI: 0.35-3.67 x 10^-9^) for WT, Δ*nucS* and *nucS*-complemented H37Rv strains, respectively (Fig. 2A and Supplementary Table 1). The small differences in mutation rates cannot be attributed to variations in drug susceptibility, as minimum inhibitory concentrations (MICs) were comparable across all strains (Supplementary Table 2), nor growth rates (Supplementary Figure 1), as growth rates were similar for H37Rv and its Δ*nucS* derivative. Taken together, these data indicate that *nucS* does not significantly influence the rate of resistance acquisition in *M. tuberculosis*.

**Figure 2.**
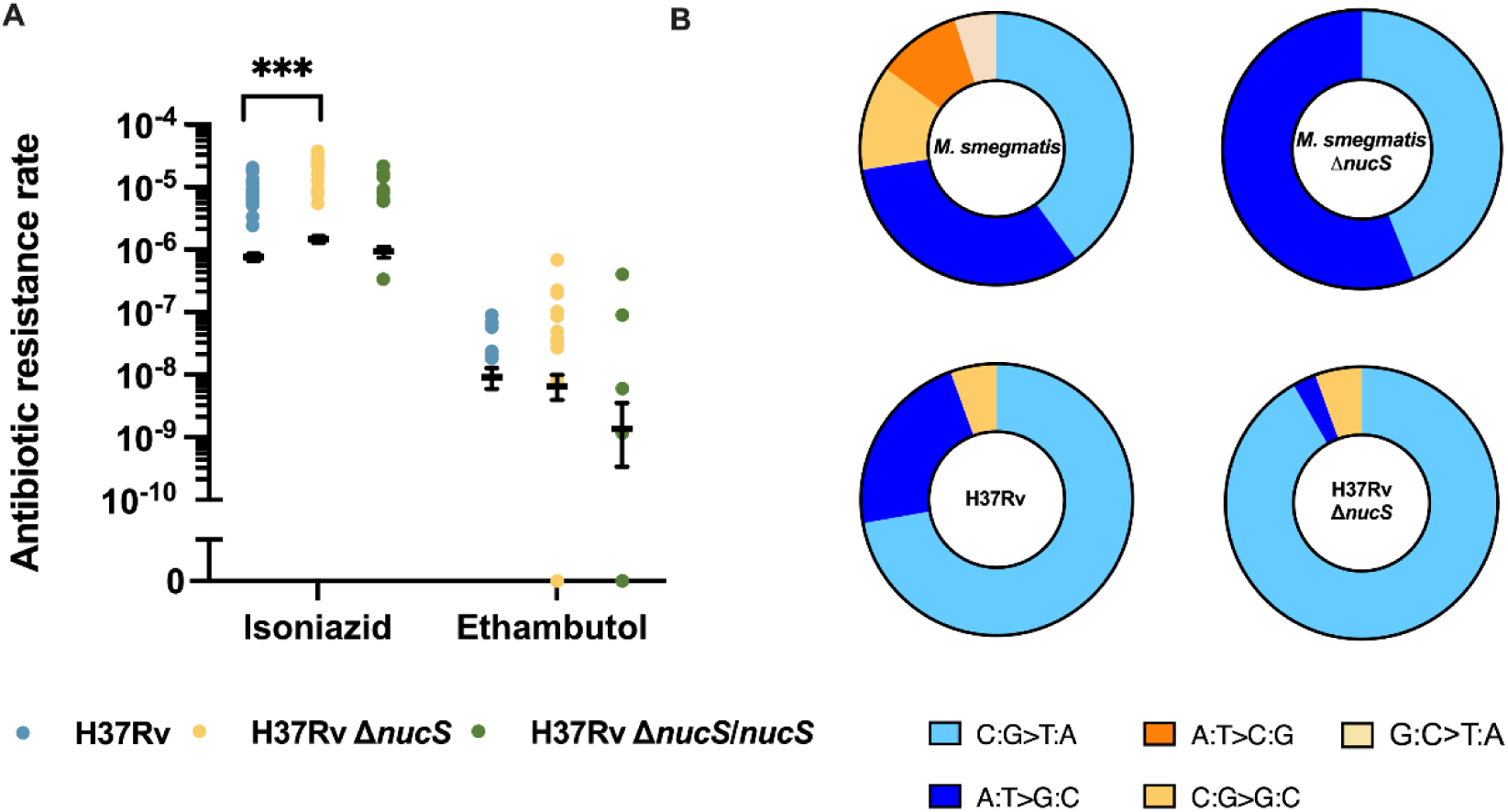
Effect of *nucS* deletion on the rate of spontaneous INH-R and EMB-R mutants in *M. tuberculosis* strains H37Rv and its derivatives (A) and mutational spectrum of strains H37Rv and its Δ*nucS* derivative, compared to those of *M. smegmatis* WT and Δ*nucS* (3) (B). Asterisks denote statistical significance (defined by nonoverlapping 99% CI). Spectra are expressed as percentage of RIF-R mutations. Transitions are shown in blue tones, transversions in orange tones. Between 1 and 5 colonies from each fluctuation test culture were pooled. Only colonies with different mutations from the same culture were considered for this study (Supplementary Table 3).

### Loss of nucS modifies the spectrum of transitions

To investigate the role of NucS in modulating base pair substitutions (BPSs) spectra in *M. tuberculosis*, we analysed RIF-R colonies by sequencing the rifampicin resistance-determining region (RRDR) of the *rpoB* gene (Supplementary Table 3). In the H37Rv WT strain, the BPS spectrum is heavily biased toward transitions, which account for 94.4% (34/36) of all mutations. Among transitions, 76.5% (26/34) were C:G > T:A, while 23.5% (8/34) were A:T > G:C. Transversions accounted for only 5.6% (2/36) (all of them C:G > G:C) (Fig. 2B and Supplementary Table 3). Important differences were found between BPSs spectra of *M. tuberculosis* H37Rv and another Mycobacteria like *M. smegmatis* mc^2^ 155, when the spectrum was analysed via mutations in the *rpoB* gene. In *M. smegmatis*, the spectrum is also biased toward transitions, although not so heavily as in H37Rv, with a 72.5%, with a 66.7% of C:G > T:A changes (3) (Fig. 2B). The transition/transversion (Tr/Tv) ratio in *M. tuberculosis* H37Rv is 17.0 (34/2) (Supplementary Table 3), significantly higher than previously reported values for other Actinobacteria, including *M. smegmatis* (2.63) (3), *C. glutamicum* (2.3) (11) and *S. ambofaciens* (1.01)(13).

Deletion of *nucS* in *M. smegmatis* led to 100% transitions (3), with a shift toward A:T > G:C (from 33.3% in WT to 56% in the Δ*nucS* strain) and the consequent decrease of C:G > T:A (from 66.7% in WT to 44% in Δ*nucS*) (Fig. 2B). However, in *M. tuberculosis* Δ*nucS*, although transitions remained predominant (94.4%, 34/36), the spectrum shifted. C:G > T:A substitutions increased from 76.5% in WT to 97.1% (33/34) in Δ*nucS* (p < 0.001; unilateral binomial test) while A:T > G:C substitutions declined from 23.5% in WT to 5.6% (2/36) (Fig. 2B and Supplementary Table 3). This suggests that, differently from *M. smegmatis*, NucS selectively prevents C:G > T:A substitutions in *M. tuberculosis*. Transversions remained unchanged, 5.6% in both Δ*nucS* and WT, and the Tr/Tv ratio in Δ*nucS* (17.0) was identical to WT. Interestingly, a high Tr/Tv ratio is a hallmark of MMR inactivation in Actinobacteria and other bacteria (see Discussion section). Therefore, the high Tr/Tv ratio in wild-type H37Rv (17.0) and its identical value upon *nucS* deletion suggest an inherently diminished MMR activity.

### Predicted effect of the H37Rv-specific R144S polymorphism on the structure of NucS

The high Tr/Tv ratio observed *M. tuberculosis* H37Rv wild-type, coupled with the lack of a significant increase in mutation rate upon *nucS* deletion and the altered transition spectrum, suggests that the NucS activity may be partially impaired in this species. Notably, sequence analysis demonstrated that the H37Rv NucS deduced sequence harbours a single amino acid substitution, arginine (R) to serine (S) at position 144 (R144S), compared to the *M. tuberculosis* consensus NucS sequence (MT 1321) from strain CDC1551(25) (Fig. 3A). Moreover, a previous work(3) proposed that this polymorphism may contribute to an increased rate of spontaneous resistant mutations, prompting us to investigate whether it could be attributed to alterations in protein conformation.

**Figure 3.**
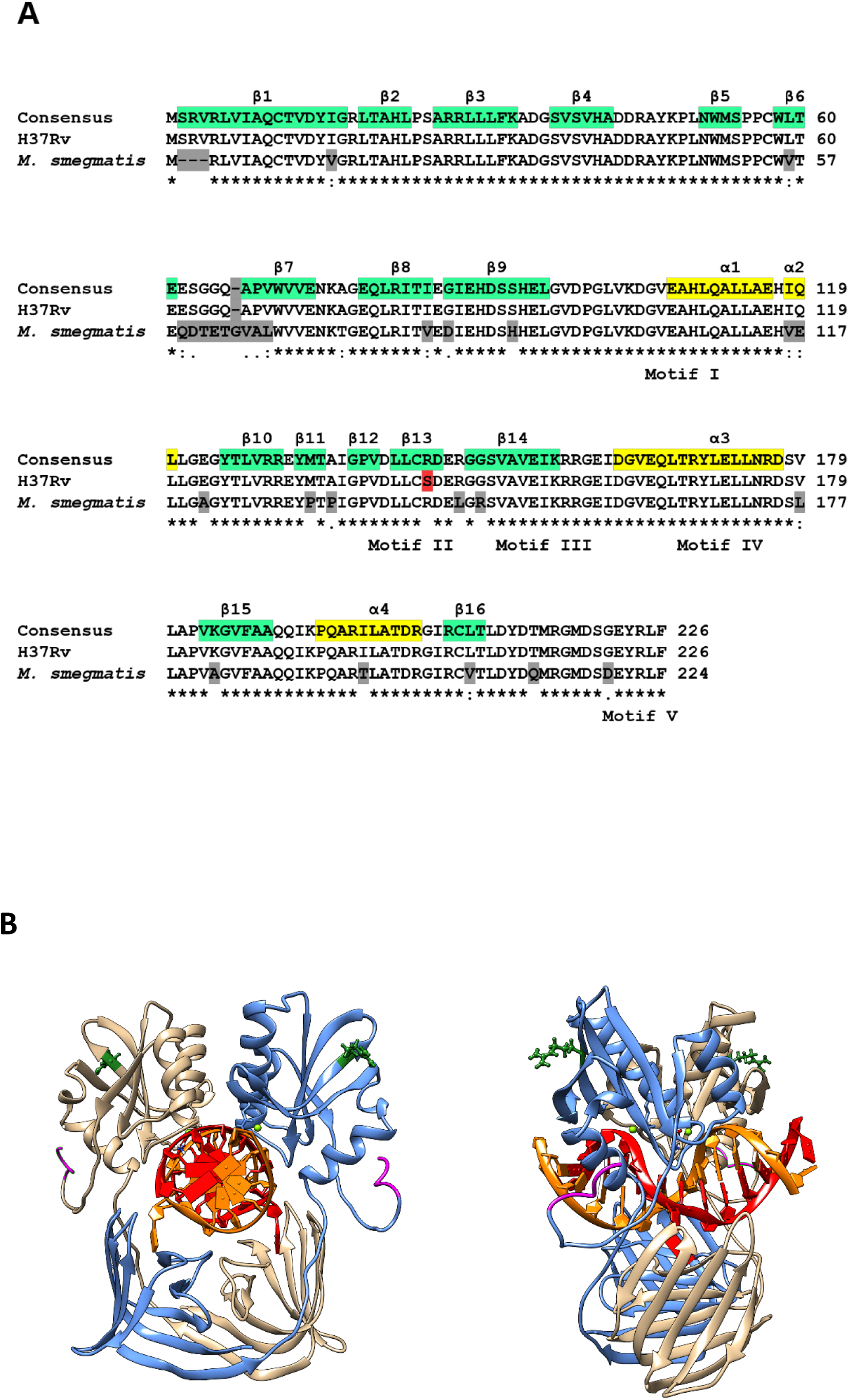
Sequence alignment and predicted structure of *M. tuberculosis* NucS-DNA complex. A) The consensus sequence of NucS from *M. tuberculosis* (NucS from strain CDC1551) is shown aligned against the sequences from *M. tuberculosis* H37Rv and *M. smegmatis* mc^2^155 using Clustal V. Differences have been highlighted in gray, except for variant R144S in *M. tuberculosis* H37Rv which has been highlighted in red to ease its identification. The degree of sequence conservation is indicated below the sequences using Clustal conventions. Sequence motifs conserved across *Archaea* by Zhang *et al.* 2019 are boxed and labelled on the bottom line (Motif V corresponds to the C-terminal PCNA binding motif). Structural features are highlighted in yellow (alpha-helix) and turquoise (beta-sheet) over the reference *M. tuberculosis* CDC1551 consensus sequence and named in the top row of the alignment. Sequence numbering is indicated on the right side for each sequence. B) The complex has been oriented and loosely colored as in Nakae et al. 2016 (7) to facilitate reference. Residue R144 is shown as ball and stick model colored in forest green, and the β-clamp-binding motif in magenta.

Firstly, to enable a comparison between wild-type and the mutant protein, it first required a structural model of *M. tuberculosis* NucS. Since no known experimental structure is currently available, we generated homology-based models of both apo (unbound) and DNA-Mg^++^-bound dimeric forms for the R144 (wild-type) and S144 (mutant) variants. These models closely align with published structures and reveal that *M. tuberculosis* NucS, while exhibiting secondary structure differences relative to archaeal homologs, retains the overall fold and conserved motifs in both conformational states (Figure 3B).

Residue 144 is located in region β-strand 13 of *M. tuberculosis* NucS C-terminal domain (corresponding to β 10 in *T. kodakarensis*), but lies outside motif V (28). Positioned on the complex outer surface and relatively distant from the C-terminal QLxxLF β-clamp binding motif (7) (EYRLF in *M. tuberculosis*), it is unlikely to directly participate in clamp binding (Fig. 3B). Its solvent-facing orientation and location outside core functional motifs further suggest it plays no direct role in DNA binding, recognition, or dimerization. However, the substitution of arginine with serine at position 144 leads to changes in both charge distribution and surface topology. The solvent-accessible surface area (SASA) of this region decreases from 358 Å^2^ with R144 to 226 Å^2^ with S144, replacing a prominent, positively charged protrusion of arginine with a flatter, dipolar and hydrophobic formed by serine (Supplementary Figure 2). These alterations indicate that residue 144 could influence interaction stability at molecular interfaces, potentially modulating recruitment or signalling within the elongation-repair complex following NucS-mediated nicking of mismatched DNA, which initiates the repair process. A STRING v11.5(29) database analysis identified several potential interaction partners, some of them related to DNA-repair, which could be influenced by the R144S substitution. They include AlkA (DNA-repair enzyme involved in the adaptive response to alkylation damage in DNA), DnaN (DNA polymerase III β-chain, β-clamp), Ogt (Methylated-DNA-protein-cysteine methyltransferase; involved in the cellular defense against the biological effects of O6-methylguanine and O4-methylthymine in DNA), RpoZ (DNA-directed RNA polymerase subunit omega), RpsT (30S ribosomal protein S20), WhiB1 and WhiB2 (transcriptional regulatory proteins). Uncharacterized proteins include Rv1312, Rv1318c (possible adenylate cyclase), Rv1322, Rv1830, Rv2413c, Rv2738c, Rv3114 (possible cytosine deaminase) (Supplementary Figure 3). These associations are meant to be specific and meaningful, i.e. proteins jointly contribute to a shared function, although it does not necessarily mean they are physically binding to each other.

### Prevalence of the R144S polymorphism among clinical isolates and its relationship with resistance

To assess whether the R144S polymorphism is disseminated among clinical isolates, we analysed a dataset comprising 46,500 *M. tuberculosis* genome sequences (30). A total of 92 unique non-synonymous polymorphisms were identified in the deduced NucS protein, including 88 missense mutations, 3 frameshifts, and 1 premature stop codon. Given that NucS consists of only 226 amino acids, this high level of variation suggests a potentially significant evolutionary role. Notably, the R144S variant was the most frequent, detected in 2,606 isolates (0.56 %).

To further investigate the potential link between R144S and antibiotic resistance acquisition in *M. tuberculosis*, we screened 14,278 genomes from the Bacterial and Viral Bioinformatics Resource Center (BV-BRC) database, which includes experimentally determined antibiotic susceptibility profiles. Consistently, R144S remained the most common NucS polymorphism, identified in 79 strains (0.55%). We conducted Chi Square and Fisher’s exact tests to assess the association between this polymorphism and resistance to 20 different antibiotics for which data were available. Among these, the NucS R144S polymorphism showed a strong statistically significant association with ethambutol resistance, but not with other antibiotics (Fig. 4A). Specifically, 1.19% (44/3,702) of EMB-R strains carried the R144S polymorphism, compared to only 0.32% (35/10,912) of EMB-susceptible strains (Fig 4B). This represents a significant enrichment (p <0.0001), suggesting a potential role for the R144S substitution in promoting ethambutol resistance.

**Figure 4.**
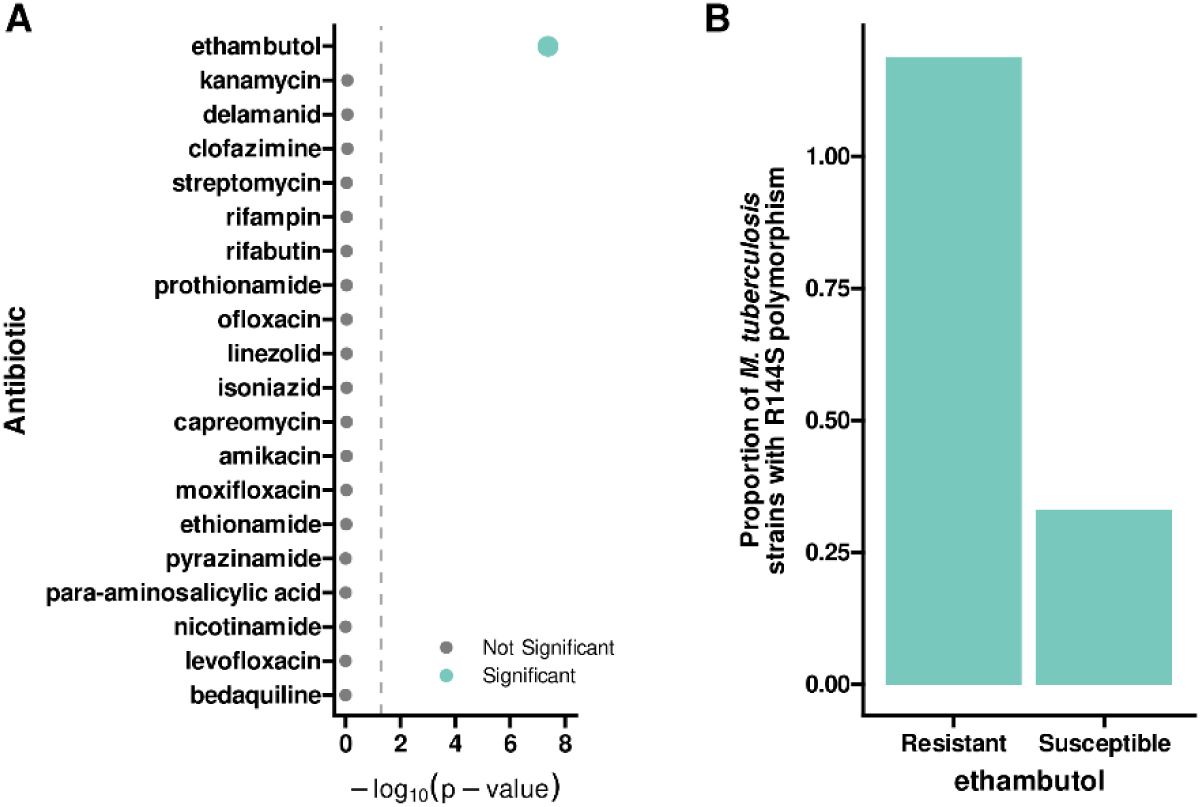
Association between the R144S NucS polymorphism and the AMR phenotype of *M. tuberculosis* strains analysed from the BV-BRC database. Statistical tests (Chi-square or Fisher’s exact test) indicated a strong association (p < 0.05 after Bonferroni-Holm adjustment) between the R144S polymorphism and resistance to ethambutol (A). Dashed grey line indicates the threshold of significance (⍺ = 0.05). The order of the antibiotics corresponds to the adjusted p values. (B) Frequency (in %) of strains carrying the R144S mutation resistant/susceptible to ethambutol (resistant, 1.19%, n = 44/3702; susceptible 0.32%, n = 35/10912).

### Effect of the change in amino acid 144 of NucS on antibiotic resistance and BPSs spectra

To directly assess the functional impact of the NucS R144S polymorphism on the acquisition of resistance mutations, we introduced the S144R (restoring the amino acid present in the consensus sequence) into mc^2^6230 and H37Rv via oligo-mediated recombineering, generating mc^2^6230-S144R and H37Rv-S144R. For the mc^2^6230-S144R strain, the RIF-R resistance mutation rate was 3.04 × 10⁻⁹ (95% CI: 2.31–3.84 ×10⁻⁹), resulting in a small 1.81-fold decrease when compared to the mc^2^6230 WT strain (Supplementary Table 1). Fluctuation tests for resistance mutation rates to RIF-R, INH-R and EMB-R in H37Rv-S144R yielded values of 3.34 × 10⁻⁹ (95% CI: 2.51–4.29 ×10⁻⁹), 6.91 × 10⁻^7^ (95% CI: 6.01–7.78 × 10⁻^7^) and 5.61 × 10⁻⁹ (95% CI: 3.75–7.83 ×10⁻⁹), respectively (Fig. 5A and Supplementary Table 1). These findings indicate that while the S144R substitution slightly alter the rate of spontaneous resistant mutants when compared to wild-type strains, it does not account for the non-mutator phenotype observed for the Δ*nucS* derivatives.

**Figure 5.**
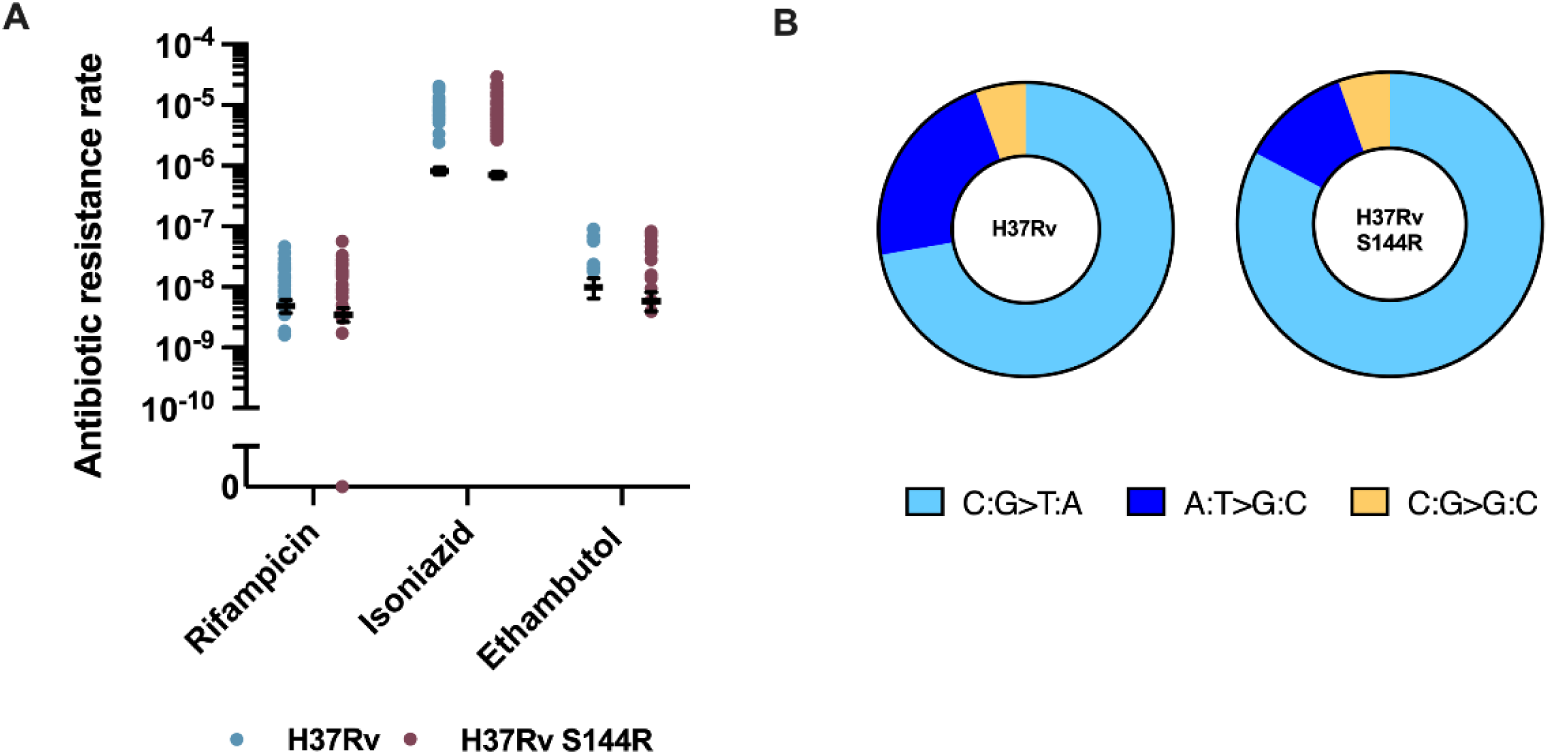
*Effect of the change in amino acid 144 of NucS on antibiotic resistance and BPSs spectrum.* Rates of spontaneous RIF-R, INH-R and EMB-R mutants in *M. tuberculosis* strains H37Rv and its S144R derivative (A) and mutational spectrum of these strains (B). Spectra are expressed as percentage of RIF-R mutations. Transitions are shown in blue tones, transversions in orange tones. Between 1 and 5 colonies from each fluctuation test culture were pooled. Only colonies with different mutations from the same culture were considered for the study (Supplementary Table 3).

However, the mutational spectrum changed upon introducing the consensus S144R sequence into NucS. While the transitions remained predominant (94.4% transitions versus 5.6% transversions) and the Tr/Tv ratio stayed identical (17), the prevalence of C:G > T:A transitions increased, although not statistically significantly, from 76.5% (WT) to 88.2% (H37Rv-S144R) (p > 0.05; unilateral binomial test), whereas A:T > G:C transitions decreased from 23.5% (WT) to 11.8% (H37Rv-S144R). These values are intermediate between the WT and the Δ*nucS* derivative (Fig. 5B and Supplementary Table 3).

Therefore, although a statistically significant relationship between the R144S substitution and ethambutol resistance was observed by analysing a genome sequence database, this association can not be attributed to an increased mutation rate to EMB-R or differences in MIC of ethambutol (Supplementary Table 2).

## Discussion

The postreplicative mismatch repair (MMR) system, which corrects mismatches that escape both base selection and 3’-5’proofreading activities of replicative DNA polymerase (2) is essential for DNA replication fidelity and the maintenance of genome stability. Traditionally, this repair was thought to be a highly conserved process mediated by members of the MutS and MutL protein families. However, the identification of NucS as a key player in genome stability in Actinobacteria and Archaea has reshaped this paradigm, establishing the existence of at least two distinct MMR pathways: the canonical MutSL system and an independently evolved NucS-dependent pathway (3). Loss or mutational inactivation of *nucS* results in a hypermutable phenotype in all Actinobacteria previously tested, underscoring its critical role in the evolution and adaptability of these industrially and clinically relevant species.

In *M. tuberculosis,* where drug resistance arises exclusively via chromosomal mutations (19), understanding the mutational landscape is, therefore, particularly relevant (20). Despite the well-established role on safeguarding genomic stability and limiting resistance acquisition in other mycobacteria, its specific function in *M. tuberculosis* remains unknown, likely due to technical challenges in generating a *nucS*-null mutant.

Here, we systematically evaluated the role of *nucS* in the acquisition of drug resistance in *M. tuberculosis*. Deletion of *nucS* has been achieved in both, the model strain H37Rv and its attenuated derivative H37Rv mc^2^6230. Unexpectedly, the deletion only marginally influenced the rate at which spontaneous RIF-R mutations appear in the mc^2^6230 strain and had no effect in H37Rv, challenging the presumed role of *nucS* as a principal guardian of genome stability and a modulator of antibiotic resistance acquisition in *M. tuberculosis*. These results contrast sharply with previous findings in other Actinobacteria, where *nucS* deletion in *M. smegmatis*(3), *S. coelicolor*(3), *S. ambofaciens*(13) and *C. glutamicum*(4, 11, 27) caused a two-orders-of-magnitude increase in RIF-R spontaneous mutation rate (Fig 1B). This non-mutator phenotype also extended to resistance acquisition to other front-line antitubercular drugs, such as INH and EMB, with only a minimal (<2-fold) increase in the INH-R mutation rate upon *nucS* deletion in H37Rv. These observations suggest the existence of additional mechanisms beyond *nucS* that control spontaneous mutagenesis in *M. tuberculosis*, warranting further investigation.

Mutational spectrum analysis provides further evidence that *nucS* functions differently in *M. tuberculosis* compared to other Actinobacteria, consistent with previous findings of G-G mismatch repair activity in H37Rv (10). Wild-type H37Rv exhibits a high transition-biased mutation spectrum (94.4% transitions; Tr/Tv ratio of 17.0), markedly higher than ratios observed in other Actinobacteria (2.63 in *M. smegmatis* (3), 2.3 in *C. glutamicum* (11), and 1.01 in *S. ambofaciens*(13)). Notably, *nucS* deletion did not affect these parameters (transitions 94.4 %; Tr/Tv ratio of 17.0), in contrast to other bacteria where MMR inactivation increases these ratios dramatically: *M. smegmatis* (38-fold)(14), *C. glutamicum* (44.6-fold)(11) and *S. ambofaciens* (36-fold)(13). Similarly, Tr/Tv ratio increases are observed in *Escherichia coli* (33-fold)(31), *Pseudomonas fluorescens* (45-fold)(32), *Deinococcus radiodurans* (9-fold)(32), *Vibrio cholerae* (29-fold)(33) and *Vibrio fisheri* (102-fold)(33). The consistently high basal Tr/Tv ratio (17.0) of the wild-type H37Rv strain and its stability upon *nucS* deletion suggest that the *nucS*-dependent MMR pathway may already be impaired, despite a baseline RIF-R mutation rate comparable to other *M. tuberculosis* strains belonging to the same lineage, such as CDC1551 (20).

A previous work proposed that naturally-occurring NucS polymorphisms may impair its function, leading to a hypermutable phenotype (3). Interestingly, the H37Rv NucS sequence carries a single R144S relative to the *M. tuberculosis* NucS consensus sequence (25). Our results suggest this variant may subtly perturb protein-protein interactions required for MMR function. Additionally, this R144S polymorphism showed a strong statistically significant association with EMB resistance in *M. tuberculosis* clinical isolates. However, fluctuation tests assessing RIF-R, INH-R and EMB-R mutation rates in engineered strains expressing the restored consensus NucS sequence (S144R) revealed minimal differences in spontaneous resistant rates. This substitution did shift the base pair substitution spectrum toward intermediate values between wild-type and *nucS*-deleted strains, although differences are not statistically significant. The mechanistic basis of the association between the R144S polymorphism and EMB resistance remains unclear, and the phenotype of the polymorphism alone does not explain the exceptionally high Tr/Tv ratio in H37Rv.

In conclusion, while *nucS* is central to maintaining genome stability and low mutation rates in Actinobacteria, its impact on mutagenesis in *M. tuberculosis* appears minimal. The persistent low mutation rate in H37Rv, even following *nucS* deletion or alteration, points to additional mechanisms governing postreplicative DNA repair and evolutionary adaptation in this pathogen. Dissecting the interplay between DNA repair pathways, mutation dynamics, and antibiotic resistance in *M. tuberculosis* will be critical to inform future therapeutic and diagnostic strategies in the ongoing battle against tuberculosis.

## Materials and methods

### Bacterial strains, culture conditions, oligonucleotides and media

The *M. tuberculosis* strains H37Rv and H37Rv mc^2^6230 (attenuated strain from the Jacob’s laboratory at Albert Einstein College of Medicine, genotypically identical to H37Rv mc^2^6030(26), except it does not contain a hygromycin resistance marker) were obtained from W. Jacobs laboratory. Strains, their mutant derivatives used in this work are listed in Supplementary Table 4. All oligonucleotides used in this work are listed in Supplementary Table 5.

*M. tuberculosis* H37Rv liquid cultures were grown on Middlebrook 7H9 (Becton Dickinson) supplemented with 0.5% glycerol, 0.05% Tween 80, 10% oleic acid, albumin, dextrose, catalase (OADC) (Becton Dickinson), and incubated at 37°C for 7-15 days. For solid media, Middlebrook 7H10 (Becton Dickinson) supplemented with 0.5% glycerol, 0.05% Tween 80, 10% OADC and 500 µg/mL amphotericin B (CAS No 1397-89-3, NeoBiotech) was used and plates were incubated at 37°C for 3-4 weeks.

When working with *M. tuberculosis* H37Rv mc^2^6230 auxotrophic strain, all media was supplemented with 24 µg/mL D-pantothenic acid hemicalcium salt (CAS No 137-08-6, Merck) and 0.2% OmniPur Casamino Acid (CAS No 65072, Calbiochem).

Media was supplemented with antibiotics when appropriate: 25 μg/mL kanamycin (Gibco), 20 μg/mL streptomycin (CAS No 3810-74-0, Merck), 50 μg/mL hygromycin (CAS No 31282-04-9, Sigma-Aldrich), 4 μg/mL rifampicin (Rifaldin, Sanofi), 2 μg/mL isoniazid (CAS No 54-85-3, Merck), 5 μg/mL thiostrepton CAS No 1393-48-2, Calbiochem), 10 μg/mL ethambutol (CAS No 1070-11-7, Merck).

Cloning was performed in *E. coli* DH5α grown in LB medium supplemented with 50 μg/mL kanamycin or 12.5 μg/mL chloramphenicol (CAS No 56-75-7, Nzytech) when appropriate.

### Construction of deletion mutants

*M. tuberculosis* H37Rv mc^2^6230 and H37Rv Δ*nucS* (Rv1321) in-frame deletion mutants were generated using the ORBIT technology(34). RecT was induced ∼16 h before transformation by addition of ATc to a final concentration of 500 ng/mL. Competent *M. tuberculosis* mc^2^6230 and H37Rv strains carrying plasmid pKM461 were transformed with 1 μg of *nucS* ORBIT oligonucleotide (Supplementary Table 5) and 200 ng of pKM464. Cells were recovered in 10 mL of 7H9 and grown for 24h at 37°C. After outgrowth, cells were plated on 7H10 plates supplemented with hygromycin. Putative deletion colonies were confirmed by PCR with oligos nucS_CDS_FW and nucS_CDS_RV; ORBIT_nucS_FW 5’-oriE Rv to verify the absence of the *nucS* gene and the 5’ junction, respectively. Additionally, the mutants were confirmed by whole genome sequencing to discard the presence of additional relevant mutations in the genome.

### Complementation of Δ*nucS* mutants

For complementation the *M. tuberculosis* Δ*nucS* mutants, the wild-type *nucS* sequence from H37Rv was cloned into the backbone of integrative vector pMV361(35). The *nucS* coding sequence and 479 bp of its upstream region were PCR amplified from genomic DNA of *M. tuberculosis* H37Rv, using oligos new-nucS_EcoRI_F and nucS_TB_HindIII_R (Supplementary Table 5). Plasmid pMV361 was PCR amplified with oligos pMV_KO_Hsp60_Eco_R and pMV_Hsp60_HindIII_F. The PCR amplification products were digested with restriction enzymes EcoRI and HindIII (New England Biolabs) and ligated using T4 ligase to eliminate the Hsp60 promoter.

The original *aph* kanamycin resistance gene from plasmid pMV361 was replaced by a chloramphenicol resistance cassette (Cam^R^). The (Cam*^R^*) was amplified by PCR from pKM464 plasmid(34) using the primers cam gene FOR and cam gene REV (Supplementary Table 5) and inserted into the plasmids carrying *nucS* previously digested with *Nhe*I-*Ase*I using the Gibson technology obtaining the pMV361 *nucS* H37Rv Cam^R^ plasmid.

Lastly, a thiostrepton resistance cassette (Tsr^R^) was amplified from pSETtsr(36) using primers tsr gene FOR and tsr gene REV (Supplementary Table 6) and inserted into the pMV361 *nucS* H37Rv Cam^R^ plasmid previously digested with *Kpn*I-*Not*I enzymes using the Gibson technology obtaining the pMV361 *nucS* H37Rv plasmid. Putative complemented mutant was obtained upon electroporation of the pMV361 *nucS* H37Rv plasmid into the *M. tuberculosis* H37Rv Δ*nucS* strain, and incubation of the plated samples on Middlebrook 7H10 agar plus thiostrepton for three weeks at 37°C. Finally, the *M. tuberculosis* H37Rv Δ*nucS* complemented mutants were analysed by PCR and Sanger sequencing.

### Identification of NucS 144 SNP in *M. tuberculosis* databases

We first interrogated a database containing 46,500 *M. tuberculosis* genome sequences previously published (30) for the presence of R144S polymorphism in the NucS deduced sequence. To analyse whether the R144S mutation of NucS of *M. tuberculosis* was associated with AMR phenotypes, the distribution of the polymorphism in a database with AMR phenotypic information was studied. The *M. tuberculosis* genomes from the strains with experimentally determined resistance phenotype data were retrieved from the Bacterial and Viral Bioinformatics Resource Center database (BV-BRC, www.bv-brc.org, accessed on 11-02-2025; n = 14,278). The presence of R144S was assessed by comparing each genome against the *nucS* gene using Snippy v.4.6.0. The *nucS* sequence from *M. tuberculosis* H37Rv, which carries the R144S mutation (RefSeq NP_215837.1), was employed as reference. The statistical significance of the association between the presence of the R144S mutation and the AMR phenotype to multiple antibiotics (n = 20) was assessed by Chi Square tests (or Fisher’s exact test if n<5 in the contingency table). P values were adjusted for multiple comparison testing by Bonferroni-Holm method.

### Construction of strains mc^2^6230-S144R and H37Rv-S144R and validation

NucS S144R mutant derivative of *M. tuberculosis* H37Rv was generated through mycobacterial oligo-mediated recombineering as previously described (37), with some modifications. In brief, 70-mer oligonucleotides were designed to correspond to the lagging strand of the replication fork at the *nucS* gene genomic region, with the desired mutation in the middle of the oligo. RecT expression was induced ∼16 h before transformation by addition of ATc to a final concentration of 500 ng/mL. 400 μl of *M. tuberculosis* H37Rv or mc^2^6230 carrying pKM461 competent cells were transformed with 2.5 µg of nucSTB_S144R oligo and 0.1 µg of rpsL_K43R_TB oligo (Supplementary Table 5). The rpsL_K43R_TB oligo generates streptomycin resistance by generating K43R SNP into the *rpsL* gene in the bacterial chromosome (37). Cells were recovered in 5 mL of 7H9 supplemented with kanamycin. After 3 days of recovery, a 1:25 dilution of the cultures was performed in 7H9 supplemented with streptomycin. This culture was grown until OD_600_ = 1 (approximately 28 days) at 37°C. And 500 μL of a 10^-6^ dilution were plated into 7H10 plates supplemented with streptomycin. The strains were cured of pKM461 plasmid by selecting individual clones growing in 7H10 plates supplemented with 3% sucrose(38). Individual colonies were picked and screened for presence of the SNPs in the *nucS* gene through PCR amplification with oligos Seq_1F_nucS_TB and Seq_1R_nucS_TB (Supplementary Table 5). The PCR amplified products were purified following the manufacturer’s instructions using QIAquick PCR Purification Kit (Qiagen) and were Sanger sequenced (StabVida) with oligo Seq_2F_nucS_TB (Supplementary Table 5) to verify the mutant genotype. Additionally, the *M. tuberculosis* H37Rv mutants were confirmed by whole genome sequencing (see below).

### Whole genome sequencing and strain validation

To check and verify the constructed strains, genomic DNA was extracted following the standard protocol for preparation of high-quality mycobacterial genomic DNA(39). The integrity of each DNA sample and the absence of RNA contamination were confirmed by DNA agarose gel electrophoresis, while its concentration and purity were measured using a NanoDrop-2000 spectrophotometer (Thermo Fisher Scientific). Genomes were sequenced through Illumina MiSeq sequencing using a MiSeq v.2 sequencing kit to obtain 250-bp paired-end reads (StabVida). The sequences were compared with those of their wild-type strain, to verify the absence of other potential mutagenic SNPs.

Paired reads were assembled against published reference genome NC_000962.3 using UGENEv52.0 (40). Raw DNAseq protocol, which encompasses read quality control with FastQC (https://www.bioinformatics.babraham.ac.uk/projects/fastqc/), quality trimming and adapter removal with CutAdapt (41), read assembly with BWA-MEM (42) and post-processing with samtools/bcftools (43) using the default parameters.

Variants with respect to the published reference genome were detected, split into SNPs and indels, and filtered (QD < 2, FS > 60, MQ < 40, SOR > 10 for SNPs, and QD < 0, FS > 200, SOR > 10 for indels) using the aligned reads and a standard best-practices protocol using GATK (44).

Identified variants were compared using bcftools to tell strain-specific from shared variants, annotated with snpEff (45) to highlight high impact and loss of function variants, and further annotated from the GTF reference annotation.

### MIC determinations

Microtiter plates to calculate Minimal Inhibitory Concentrations (MICs) were prepared by serially diluting 4X stocks of rifampicin (4 µg/mL – 0.008 µg/mL) and isoniazid (4 µg/mL – 0.008 µg/mL) with 7H9 on sterile, clear, round bottomed, 96-well microplates (Thermo Scientific™ Nunc MicroWell). After outgrowth on 7H9, 1:200 cell suspensions were prepared for *M. tuberculosis* mc^2^6230, H37Rv and their *nucS* mutant derivatives. Plates were inoculated with 100 µL of the cell suspension. To prevent evaporation, 200 μL distilled H_2_O was added to the outer perimeter wells. Each plate included a negative control and a growth control. Plates were covered with their lids, placed in individual plastic bags and incubated at 37°C for 6 days. After incubation, 30 μL of 100 μg/mL resazurin sodium salt solution (CAS-No: R7017-1G, Sigma-Aldrich) were added to each well and further incubated for 48 hours at 37°C. MICs were assessed using a microplate reader with 555 nm excitation and 590 nm emission (FLUOstar Omega). Each MIC had 3 technical replicates. In the case of ethambutol, MIC was evaluated in solid medium. Cultures replicates of H37Rv and its mutant derivatives were grown until OD_600_=1. Lastly, 5 μL of the different strains were added to plates containing different concentrations of ethambutol ( 0, 1, 2, 4, 8 and 16 μg/ml) and growth was checked after three weeks.

### Bacterial growth

Plasmid pYUB3054, is a plasmid carrying GenL, a *M. tuberculosis* codon-optimized nanoluciferase (nluc*_M. tuberculosis_*)(46) amplified from the Addgene plasmid #85200, the integration system of phage tweety(47) and an *aph* kanamycin resistance cassette.

Plasmid pYUB3054 was transformed into *M. tuberculosis* H37Rv and its *nucS* mutant derivatives. Bacterial growth was evaluated by growth curve assays of *M. tuberculosis* H37Rv and its *nucS* mutant derivatives and using the correlation between relative luminescent units (RLUs) and viable cells. Four independent 10 mL cultures starting at OD_600_ 0.05 were incubated at 37°C in static condition and nanoluciferase activity was measured every 24 hours for a total of 18 days. RLUs were evaluated on white, 96-well microplates (Ref 655073, Greiner Bio-One) and measuring nanoluciferase activity. The Nano-Glo Luciferase Assay System (Promega) was used following manufacturer’s instructions. Briefly, Nano-Glo substrate was diluted 1:50 in Nano-Glo buffer and mixed 1:1 (vol/vol) with the samples. After addition of substrate, plates were covered with adhesive, optically clear plate sealer (Microseal B PCR plate-sealing film, adhesive, optical no. MSB1001; Bio-Rad) and decontaminated, and luminescence was read immediately on a FLUOstar Omega spectophotometer.

Growth curves were compared using a Baranyi model fitted using R (R Core Team (2024) R: A Language and Environment for Statistical Computing. R Foundation for Statistical Computing, Vienna, Austria; https://www.r-project.org) package Growth rates and, to avoid biases due to experimental conditions, μ-max (maximum growth rate) values were checked for normality with Shapiro-Wilk test and for homocedasticity with Bartlett’s test, and differences were compared using ANOVA followed by Tukey *post hoc* tests.

### Estimation of mutation rates

Mutation rates of *M. tuberculosis* H37Rv mc^2^6230, H37Rv and their *nucS* mutant derivatives to each antibiotic were determined by fluctuation tests and validated by up to 4 different experiments each. Briefly, a starting culture of each strain was diluted 1:1,000 to generate 8 independent liquid cultures in Middlebrook 7H9 broth and incubated 15 days at 37°C. All grown cultures (10^9^ to 10^10^ cells/mL) and serial dilutions were plated on Middlebrook 7H10 agar plates without antibiotic (for estimation of viable cells) and with antibiotic (for drug-resistant cells) and incubated for 28 days at 37°C. After this incubation, CFUs were counted to obtain the total number of viable cells (Nt) and mutant cells in the cultures.

To estimate mutation rate (μ), data was pooled from multiple fluctuation test experiments as previously described by Zheng (2023)(48). Briefly, the mutation rate (μ) was recasted as a function of the parameter β, defined by μ = e^β^. In this way, the maximum likelihood estimate (MLE) of β was found using the optimize function in R, and the 95% confidence intervals (CIs) for β were constructed based on the likelihood ratio principle using R function uniroot.

The 99% confidence intervals were where constructed as previously described and where used exclusively to compare mutation rates at the 0.001 significance levels (49). This approach was chosen to address the non-standard problem of pooling data from different fluctuation test experiments. As described by Zheng(50), using overlapping confidence intervals is particularly suitable for comparing mutation rates when experiments involve varying terminal population sizes, ensuring reliable statistical conclusions without simplifying assumptions.

### Mutational spectra

The mutational spectra of *M. tuberculosis* strains H37Rv and its mutant *nucS* derivatives were characterized by picking individual resistant colonies from RIF plates. Between 1 and 5 colonies per plate were isolated, and colonies from all the fluctuation tests were pooled, allowing for more than 40 colonies per strain. To avoid estimating mutations from the same original mutant, only different mutations from the same culture were considered for the study of mutational spectrum.

For resistance to rifampicin, the rifampicin resistance-determining region in the *rpoB* gene was PCR amplified using a pair or primers RifRRDR-H37Rv Fw and RifRRDR-H37Rv Rv (Supplementary Table 3). PCR products were sequenced using oligo RifRRDR-H37RvFW and compared to the reference gene sequences to identify the type of mutation in each RIF-resistant isolate.

Tr/Tv rates were compared using a Log likelihood ratio test and differences in spectra were compared using a proportions test with Yatés continuity correction.

### Protein structure prediction

We used the protein sequence of *M. tuberculosis* strain CD1551 NucS from entry P59979.1 to build a full-sequence 3D structural model using I-TASSER (51). To model the dimer, we next used as reference the published structures 5GKE, 5GKF, 5GKG, 5GKH, 5GKI and 5GKJ(7) from PDB(52). After selecting the best matching protein model, we superposed two CD1551 subunits over the available dimers, both in the apo and DNA-bound forms, then optimized the structures using the Amber force field, checked the results for conflicts and selected the best models using UCSF Chimera(53). The DNA-Mg^++^-bound dimer was then used to model the structure of the S144 variant using Modeller(54) with a very-large refinement, the predicted structure was subjected to additional minimization using Amber and inspected in UCSF Chimera.

## Acknowledgements

Q. Zheng for his kind advice on statistics for mutation rates comparisons, Jeremy Rock for kindly allowing access to the Rockefeller *M. tuberculosis* dataset. Alfredo Castañeda and Esmeralda Cebrián for preliminary experiments with deletion mutants and Amparo Errandonea for helping with figures.

## Data availability

All data supporting the findings of this study are available within the paper and its Supplementary Information. Row WGS data were deposited into the NCBI Sequence Read Archive (SRA) under accession code PRJNA1277151 and will be publicly available after publication.

## Supplementary Material

**Supplementary Table 1.** Mutation rates (µ, mutations/cell/generation) calculated from fluctuation tests for RIF-R, INH-R and EMB-R. Number of replicates (n) per strain and antibiotic, 95% confidence interval (CI) and fold increase compared to each respective wild-type strain (set to 1.00). Three stars: p ≤ 0.001.

| Strain | RIFAMPICIN |  |  |  | ISONIAZID |  |  |  | ETHAMBUTOL |  |  |  |
| --- | --- | --- | --- | --- | --- | --- | --- | --- | --- | --- | --- | --- |
| | n | $\mu$ | CI 95% | Fold increase | n | $\mu$ | CI 95% | Fold increase | n | $\mu$ | CI 95% | Fold increase |
| mc<sup>2</sup>6230 | 51 | $5.49 \times 10^{-9}$ | $(4.65-6.37) \times 10^{-9}$ | 1 | | | | | | | | |
| mc<sup>2</sup>6230 ΔnucS | 26 | $1.55 \times 10^{-8}$ | $(1.24-1.89) \times 10^{-8}$ | 2.83*** | | | | | | | | |
| mc<sup>2</sup>6230 S144R | 32 | $3.04 \times 10^{-9}$ | $(2.31-3.84) \times 10^{-9}$ | 0.55*** | | | | | | | | |
| H37Rv | 32 | $4.66 \times 10^{-9}$ | $(3.58-5.86) \times 10^{-9}$ | 1 | 23 | $8.09 \times 10^{-7}$ | $(6.95-9.20) \times 10^{-7}$ | 1 | 8 | $9.57 \times 10^{-9}$ | $(6.20-13.30) \times 10^{-9}$ | 1 |
| H37Rv ΔnucS | 16 | $5.88 \times 10^{-9}$ | $(4.18-7.79) \times 10^{-9}$ | 1.26 | 20 | $1.57 \times 10^{-6}$ | $(1.37-1.76) \times 10^{-6}$ | 1.94 *** | 16 | $6.88 \times 10^{-9}$ | $(4.10-10.40) \times 10^{-9}$ | 0.72 |
| H37Rv ΔnucS/nucS | 8 | $3.33 \times 10^{-9}$ | $(1.82-5.29) \times 10^{-9}$ | 0.71 | 8 | $9.96 \times 10^{-7}$ | $(7.89-11.9 \times 10^{-7})$ | 1.23 | 8 | $1.42 \times 10^{-9}$ | $(0.35-3.67) \times 10^{-9}$ | 0.15 *** |
| H37Rv S144R | 32 | $3.34 \times 10^{-9}$ | $(2.51-4.29) \times 10^{-9}$ | 0.72 | 24 | $6.91 \times 10^{-7}$ | $(6.01-7.78) \times 10^{-7}$ | 0.85 | 16 | $5.61 \times 10^{-9}$ | $(3.75-7.83) \times 10^{-9}$ | 0.59 |

**Supplementary Table 2.**
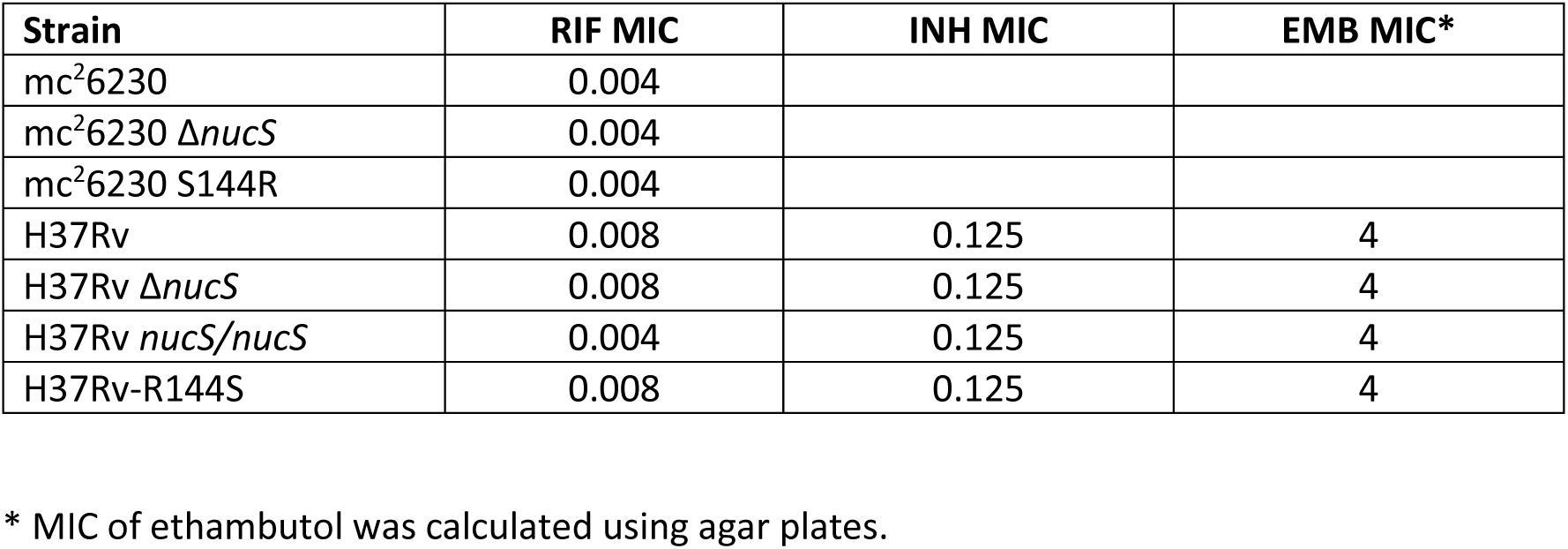
MICs of antibiotics for strains.

| Strain | RIF MIC | INH MIC | EMB MIC* |
| --- | --- | --- | --- |
| mc<sup>2</sup>6230 | 0.004 |  |  |
| mc<sup>2</sup>6230 ΔnucS | 0.004 | | |
| mc<sup>2</sup>6230 S144R | 0.004 |  |  |
| H37Rv | 0.008 | 0.125 | 4 |
| H37Rv ΔnucS | 0.008 | 0.125 | 4 |
| H37Rv nucS/nucS | 0.004 | 0.125 | 4 |
| H37Rv-R144S | 0.008 | 0.125 | 4 |

\* MIC of ethambutol was calculated using agar plates.

**Supplementary Table 3.** Mutational spectrum of strains H37Rv wild-type and its derivatives H37Rv-Δ*nucS* and H37Rv-S144R as analysed from sequences of *rpoB* of RIF-R mutants. 30 to 51 colonies from each strain were analysed yet only those containing different mutations from the same culture were considered for the study of mutational spectrum.

| Position | Codon change | Amino acid change | H37Rv | H37Rv ΔnucS | H37Rv-S144R |
| --- | --- | --- | --- | --- | --- |
| C 1322 G (Tv) | TCG→TGG | Ser 441 Trp | 1 |  |  |
| C 1322 T (Tr) | TCG→TTG | Ser 441 Leu | 1 |  |  |
| C 1333 T (Tr) | CAC→TAC | Hys 445 Tyr | 8 | 16 | 18 |
| C 1333 G (Tv) | CAC→GAC | Hys 445 Asp | 1 | 1 | 1 |
| A 1334 G (Tr) | CAC→CGC | Hys 445 Arg | 8 | 1 | 4 |
| A 1334 C (Tv) | CAC→CCC | Hys 445 Pro |  |  |  |
| A 1336 C (Tv) | AAG→CAG | Lys 446 Gln |  |  |  |
| C 1349 T (Tr) | TCG → TTG | Ser 450 Leu | 17 | 17 | 12 |
| C 1349 G (Tv) | TCG → TGG | Ser 450 Trp |  | 1 | 1 |
| Colonies analysed |  |  | 44 | 63 | 42 |
| **TOTAL BPS** |  |  | **36** | **36** | **36** |
| **% Transitions/BPS** |  |  | **94.4% (34/36)** | **94.4% (34/36)** | **94.4% (34/36)** |
| % C:G>T:A |  |  | 76.5% (26/34) | 97.1% (33/34) | 88.2% (30/34) |
| % A:T>G:C |  |  | 23.5% (8/34) | 2.9% (1/34) | 11.8% (4/34) |
| **% Transversions/BPS** |  |  | **5.6% (2/36)** | **5.6% (2/36)** | **5.6% (2/36)** |
| % C:G>G:C |  |  | 100% (2/2) | 100% (2/2) | 100% (2/2) |
| % A:T>C:G |  |  | 0% | 0% | 0% |
| Tr/Tv ratio |  |  | **17** | **17** | **17** |

**Supplementary Table 4.** Bacterial strains and their mutant derivatives.

| Strain name | Genotype | Description | Markers | Source |
| --- | --- | --- | --- | --- |
| H37Rv mc<sup>2</sup>6230 | ΔRD1 ΔpanCD | Attenuated strain; auxotrophic for pantothenate | None | W. R. Jacobs |
| H37Rv | wild type |  | None | W. R. Jacobs |
| H37Rv mc<sup>2</sup>6230 ΔnucS | ΔnucS::pKM464 | mc<sup>2</sup>6230 deleted of Rv1321 by ORBIT | Hyg<sup>R</sup> | This study |
| H37Rv ΔnucS | ΔnucS::pKM464 | H37Rv deleted of Rv1321 by ORBIT | Hyg<sup>R</sup> | This study |
| H37Rv ΔnucS / nucS | H37Rv ΔnucS complemented with Rv1321 gene | H37Rv ΔnucS; pMV361 nucS H37Rv integrated at attB site (tRNA<sup>Gly</sup>) | Hyg<sup>R</sup>Cam<sup>R</sup> Tsr<sup>R</sup> | This study |
| H37Rv mc<sup>2</sup>6230-S144R | nucS S144R | Mutant generated by oligo-mediated recombineering | None | This study |
| H37Rv-S144R | nucS S144R | Mutant generated by oligo-mediated recombineering | None | This study |

**Supplementary table 5.**
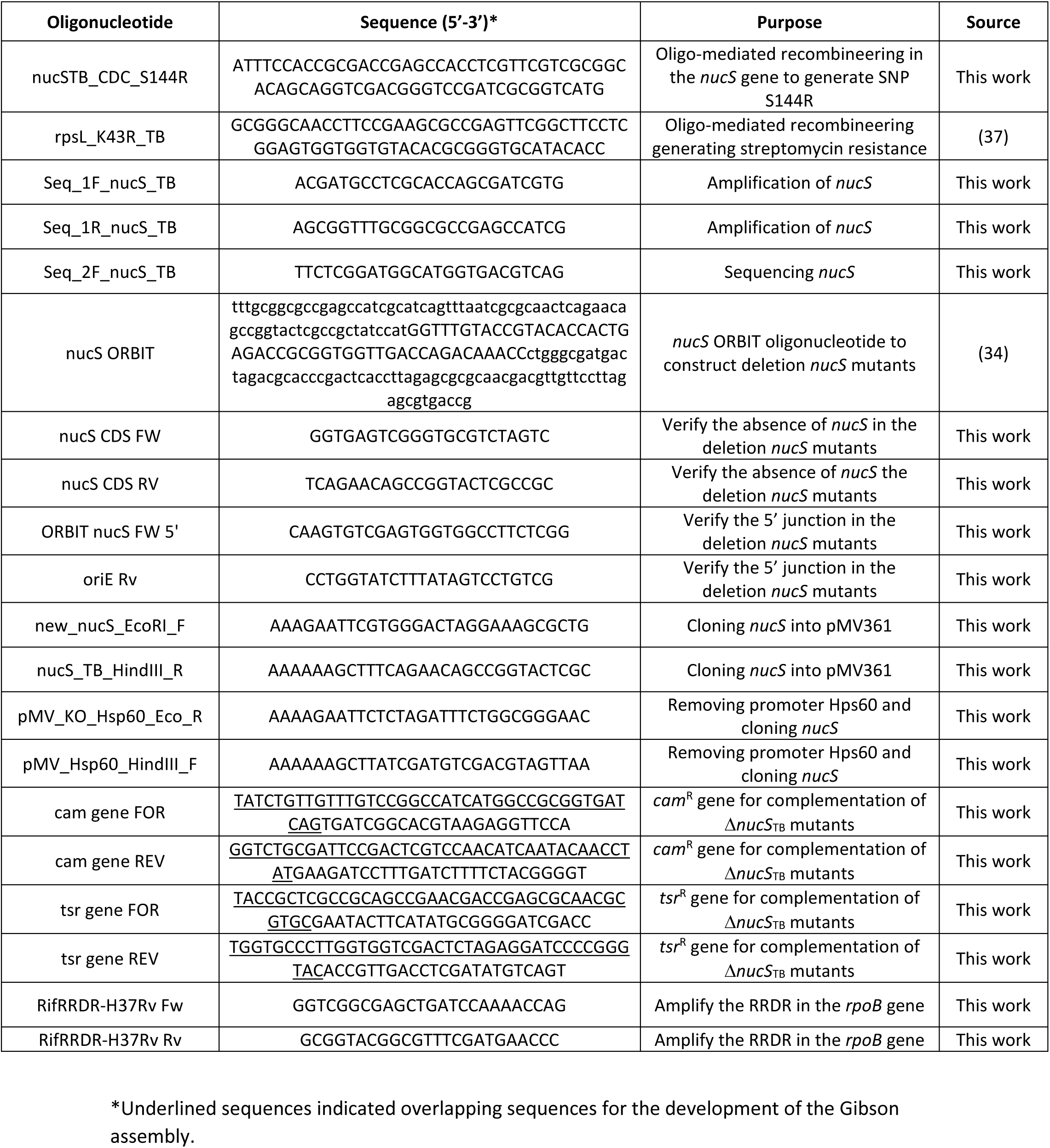
Oligonucleotides used in this work.

## Supplementary Figures

**Supplementary Figure 1.**
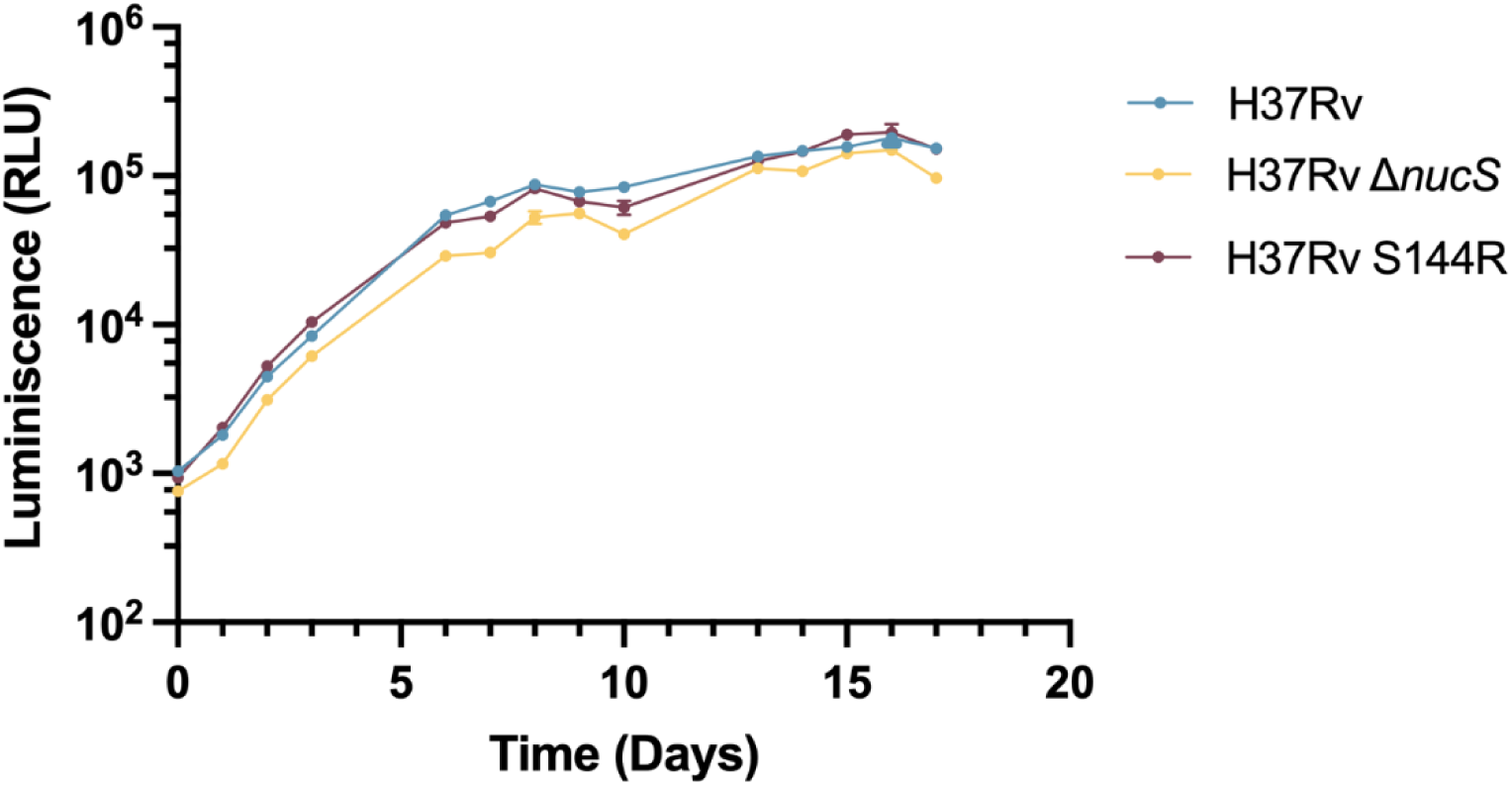
Growth curves of *M. tuberculosis* H37Rv, H37Rv Δ*nucS* and H37Rv-S144R measured by nanoluciferase activity every 24 hours for a total of 18 days. Each point is the median value from four independent cultures. Error bars represent the SEM. Differences in maximum growth rates are not statistically significant (ANOVA followed by Tukey *post hoc* tests).

**Supplementary Figure 2:**
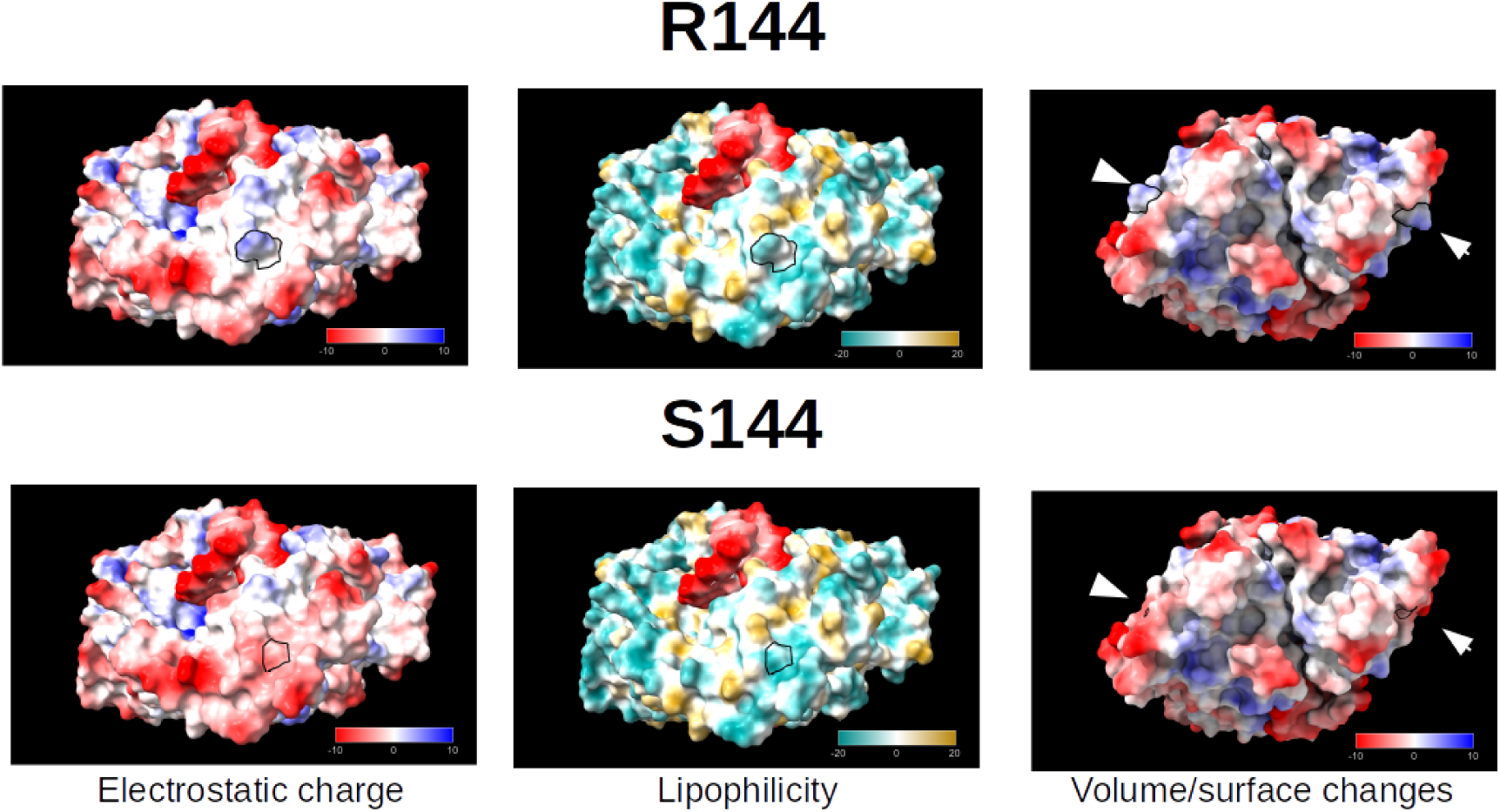
Surface structural changes associated with arginine or serine in position 144 of NucS. The left and center columns show electrostatic charge and lipophilicity respectively, mapped on the protein-DNA complex surface, and oriented so that residue 144 is approximately centered (and delineated in black) to better highlight the changes they induce in their surrounding region. dsDNA stands out in red in the central panes. The right column shows electrostatic charge, but has been oriented to show both 144 residues-delineated in black and indicated by arrows- to highlight the volume changes in the protein surface. Substitution of R by S removes its positive charge and allows exposition of additional negative charge in its surrounding environment, induces lipophilicity changes, and causes a noticeable reduction in the exposed volume and surface area.

**Supplementary Figure 3:**
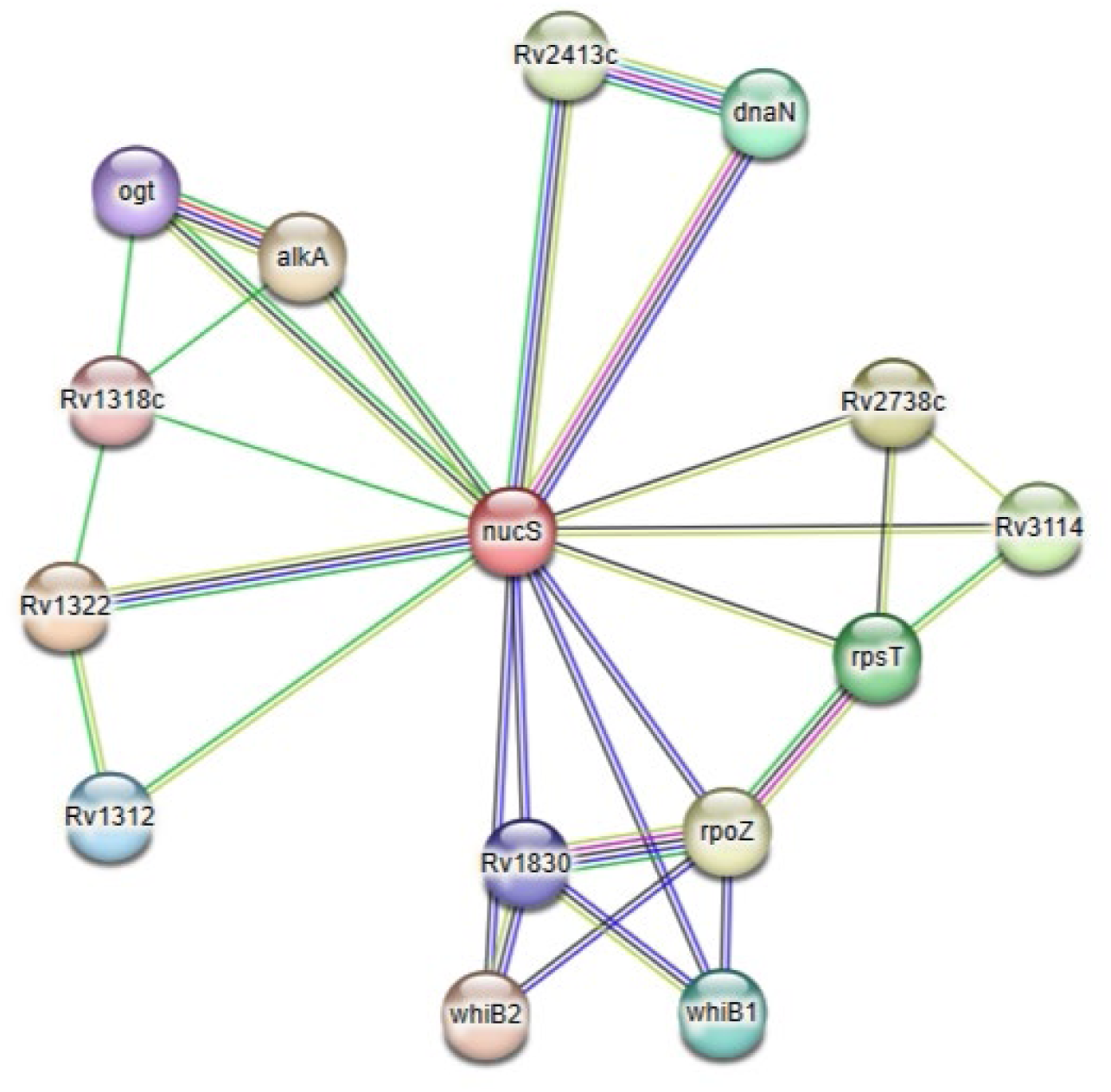
Predicted *M. tuberculosis* H37Rv *nucS* interaction network computed using STRING v 12.0 (https://doi.org/10.5167/uzh-229577). The network shows only predicted direct interactions, and was calculated using a high (score > 0.7) confidence level. Associations are meant to be specific and meaningful, i.e. proteins jointly contribute to a shared function; this does not necessarily mean they are physically binding to each other. The nodes are labelled with gene names and have been colored according to their grouping using k-means for four clusters. Edges are colored by interaction evidence:

---

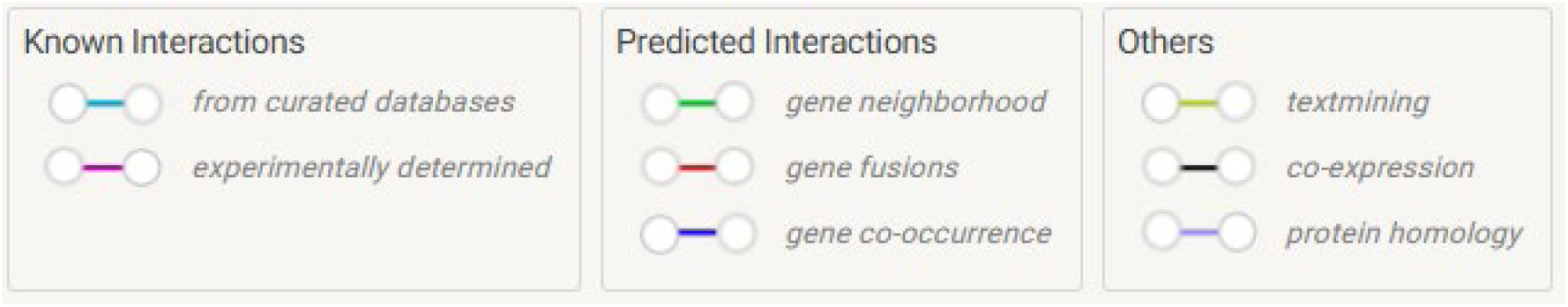

